# Modelling SARS-CoV-2 spike-protein mutation effects on ACE2 binding

**DOI:** 10.1101/2022.08.25.505249

**Authors:** Shivani Thakur, Rajaneesh Kumar Verma, Kasper Planeta Kepp, Rukmankesh Mehra

## Abstract

The binding affinity of the SARS-CoV-2 spike (S)-protein ΔΔG_bind_ to the human membrane protein ACE2 is critical for virus function and evolution. Computational structure-based screening of new S-protein mutations for ACE2 binding lends promise to rationalize virus function directly from protein structure and ideally aid early detection of potentially concerning variants. We used a computational protocol based on cryo-electron microscopy structures of the S-protein to estimate the ACE2-binding that gave good trend agreement with experimental ACE2 affinities. We then expanded predictions to all possible S-protein mutations in 21 different S-protein-ACE2 complexes (400,000 ΔΔG_bind_ data points in total), using mutation group comparisons to reduce systematic errors. We show that mutations that have arisen in major variants as a group maintain ACE2 affinity significantly more than random mutations in the total protein, at the interface, and at evolvable sites, with differences between variant mutations being small relative to these effects. Omicron mutations as a group had a modest change in binding affinity compared to mutations in other major variants. The single-mutation effects are consistent with ACE2 binding being optimized and maintained in omicron, despite increased importance of other selection pressures (antigenic drift). As epistasis, glycosylation and in vivo conditions will modulate these effects, computational predictive SARS-CoV-2 evolution remains far from achieved, but the feasibility of large-scale computation is substantially aided by using many structures and comparison of mutation groups rather than single mutation effects, which are very uncertain. Our results demonstrate substantial challenges but indicate ways to improve the quality of computer models for assessing SARS-CoV-2 mutation effects.

## 1. Introduction

Since early 2020, the severe acute respiratory syndrome coronavirus 2 (SARS-CoV-2) has caused disease across the planet, with casualties counted in many millions [1–3]. Major research efforts are directed towards understanding the structure-function relationships of the virus spike-protein (S-protein), which infects human host cells via binding to human angiotensin-converting enzyme 2 (ACE2) [4–7]. While this fusion process is central to infection, it can be partly prevented by immune recognition and neutralization of S-protein epitopes, which is the rationale behind many important vaccines.[8,9] For this reason, antigenic drift leading to new variants that evade vaccine-induced antibodies is a central concern [10–13]. The highly mutated omicron variant illustrates this principle perfectly, with its associated large transmission in already vaccinated people since December 2021 [14,15].

The S-protein on the surface of the virus features a dynamic, metastable pre-fusion state that converts into an open state upon ACE2 binding, with the receptor binding domain (RBD) in an upwards conformation [16–20]. The abilities to bind to ACE2 and evade antibodies depend on this structure and its conformational variations [21]. The emergence of new variants that maintain or improve ACE2 binding will probably remain important for many years to come [22]. It is of major interest, both scientifically and for pandemic surveillance via the prediction of future evolution potential of the virus, to understand ACE2 binding at the molecular level of the amino acids.

Some optimization of ACE2 binding occurred via the D614G mutation which rapidly became dominant in 2020 [23], and ACE2 binding was most likely further improved in subsequent variants alpha, beta, and gamma, via e.g., N501Y and S477N and E484K mutations [24]. The mutation L452R may play a key role in facilitating evolution [25], possibly by modulating changes in electrostatic interactions via positive charges with the predominantly negatively charged ACE2 surface [21,26,27]. There is some disagreement on whether omicron has less [28] or similar [29] binding to ACE2 compared to delta. This difference is important because weaker binding could not relate to clinical presentation (upper airway infection) but could also indicate that antigenic drift dominates ACE2 binding in the fitness function of the virus.

Computational models are necessary to understand mutation effects mechanistically and could ideally enable complete early screening of mutations of potential concern many weeks before epidemiological and clinical data become more established. Many mutations may occur outside the RBD (for which many mutations have been studied experimentally [30]), and computer models could ideally account for features not in experimental screening assays (e.g. change in conditions and protein state) and cover the full sequence space, ideally to monitor the pandemic potential of future mutations. The publication of hundreds of cryo-electron microscopy structures of the S-protein, facilitated by healthy competition between many research groups, makes a structure-guided computation much more feasible than for many other proteins [21]. However, there are major technical challenges to modeling such mutation effects, relating to the accuracy of the models and the biases they carry from training on biased mutation data sets [31–33], the use of a static experimental wild-type structure to extrapolate the impact of mutations [34,35], the quality of this structure and the sensitivity of results to changes in used structure [36,37], and the biological relevance of the experimental cryo-EM structures to in vivo conditions, e.g. effects of chemical conditions, solvent, and protein modifications [38].

In this paper, we use a computational protocol to describe the effect of all possible mutations in the S-protein on ACE2 binding, after benchmarking it against RBD mutation effects, where experimental data are available. We discuss uncertainties in the methods and show how results depend on the choice of cryo-EM structure used and that mutation group comparisons increase the significance of such computational screening by reducing the uncertainty present for individual mutation effects. Our results demonstrate substantial challenges but also indicate necessary steps forward towards predictive computer models of SARS-CoV-2 evolution.

## 2. Methods

### 2.1 Protein structural data

As one of the new approaches in this work, made feasible by the rich structural data of the S-protein, we study the impact of using different structures on the computed ACE2 binding effects. The S-protein displays large conformational changes in vivo [17,19]. Computational studies published so far used only a single cryo-EM structure for each state of interest (prefusion protein, antibody-bound, ACE2-complex) [39–44]. However, the heterogeneity of experimental structures deduced by cryo-electron microscopy (cryo-EM) may impact the results of structure-based computational modeling [21], i.e., results reported from one structure may not be statistically representative of the ensemble of relevant conformation states, perhaps even for the same state. To understand these methodological dependencies we studied 21 different S-protein structures in complex with ACE2 deposited in the Protein Data Bank (PDB) [45] as a sensitivity test.

We expect the structures to have good backbone resolution and accuracy at secondary and tertiary structure level [46]. However, cryo-temperature is likely to freeze out some conformations and causes cryo-contraction of the overall protein [38]. This produces some variation relative to the physiological state, which is also heavily glycosylated in ways not accounted for by the experimental structures. Plausibly, sites shielded by glycans will not change interaction with ACE2 as much upon mutation as predicted from non-glycosylated proteins. The realism of chemical composition, notably protein modification, may also modify results both from computational studies and experiments. As a rationale for our protocol, we expect the average of the identified 21 structures to provide a reasonable consensus and sensitivity estimate for the variation in molecular ACE2-S-protein interactions seen structurally.

The structures 7A94, 7A95, 7A96, 7A97, and 7A98 by Benton et al. [47], 7CT5 (Guo et al.) [48], 7DF4 (Xu et al., 2021) [49], 7DX5, 7DX6, 7DX7, 7DX8, and 7DX9 (Yan et al., 2021) [50], 7KJ2, 7KJ3, and 7KJ4 (Xiao et al., 2021) [51], and 7KMS, 7KMZ, 7KNB, 7KNE, 7KNH, 7KNI (Zhou et al., 2020) [52] were selected among original variant structures, to make all mutations into these structures feasible and comparable, with the requirement of nearly complete trimer structures (number of residues N > 1000) [21]. The number of residues (N), percentage outliers in Ramachandran plot (% Outliers) and resolution are summarized in **Table 1**. The resolution varies from 2.9 to 5.4 Å (**Table 1**). In these resolution ranges the main-chain conformations are largely well-resolved but the accuracy of the side-chain conformations should not be over-emphasized at the given level of cryo-EM structure resolutions [46], and multiple structures may be beneficial.

**Table 1.**
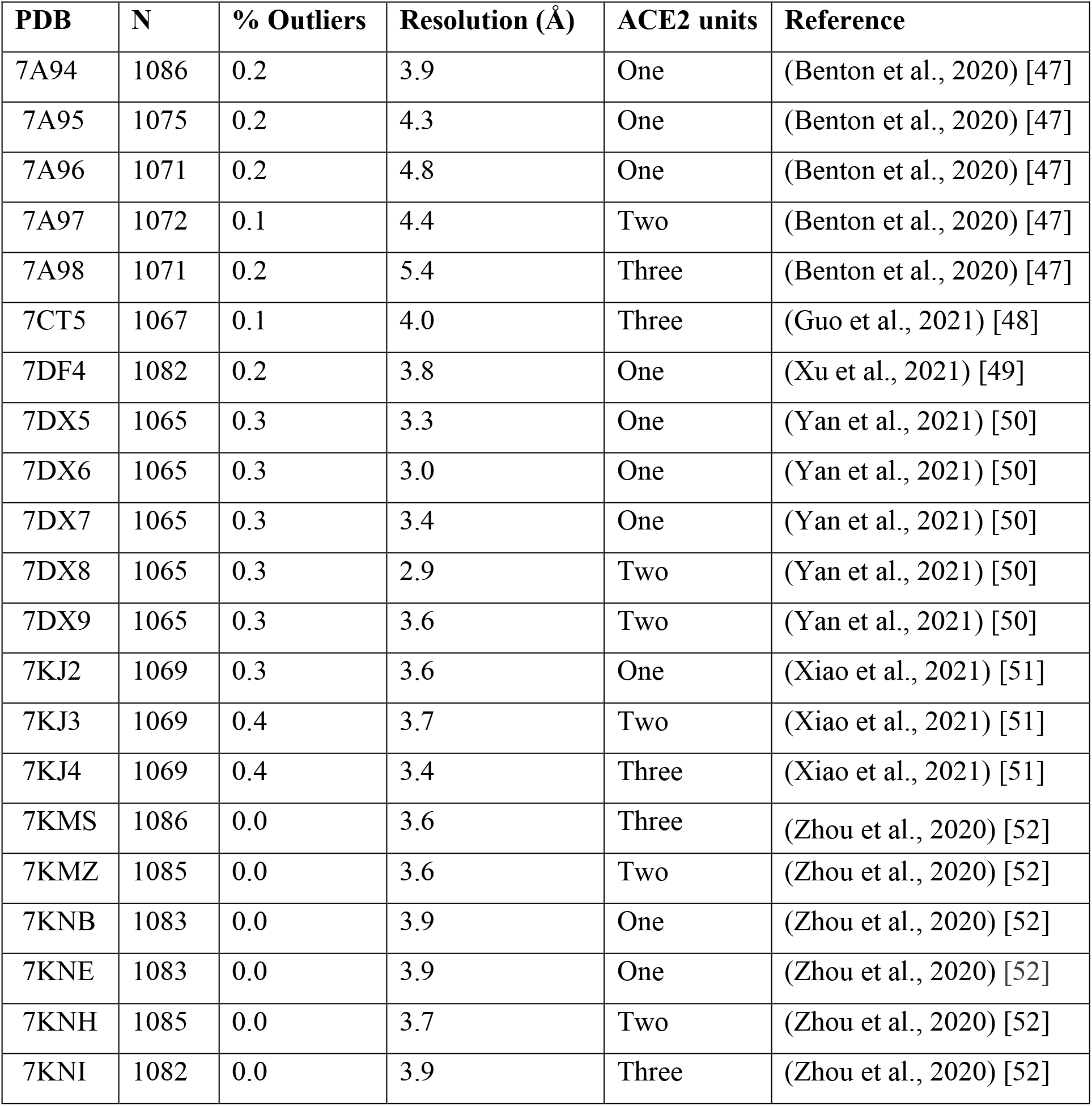
S-protein-ACE2 complexes used in the present study.

| PDB | N | % Outliers | Resolution (Å) | ACE2 units | Reference |
| --- | --- | --- | --- | --- | --- |
| 7A94 | 1086 | 0.2 | 3.9 | One | (Benton et al., 2020) [47] |
| 7A95 | 1075 | 0.2 | 4.3 | One | (Benton et al., 2020) [47] |
| 7A96 | 1071 | 0.2 | 4.8 | One | (Benton et al., 2020) [47] |
| 7A97 | 1072 | 0.1 | 4.4 | Two | (Benton et al., 2020) [47] |
| 7A98 | 1071 | 0.2 | 5.4 | Three | (Benton et al., 2020) [47] |
| 7CT5 | 1067 | 0.1 | 4.0 | Three | (Guo et al., 2021) [48] |
| 7DF4 | 1082 | 0.2 | 3.8 | One | (Xu et al., 2021) [49] |
| 7DX5 | 1065 | 0.3 | 3.3 | One | (Yan et al., 2021) [50] |
| 7DX6 | 1065 | 0.3 | 3.0 | One | (Yan et al., 2021) [50] |
| 7DX7 | 1065 | 0.3 | 3.4 | One | (Yan et al., 2021) [50] |
| 7DX8 | 1065 | 0.3 | 2.9 | Two | (Yan et al., 2021) [50] |
| 7DX9 | 1065 | 0.3 | 3.6 | Two | (Yan et al., 2021) [50] |
| 7KJ2 | 1069 | 0.3 | 3.6 | One | (Xiao et al., 2021) [51] |
| 7KJ3 | 1069 | 0.4 | 3.7 | Two | (Xiao et al., 2021) [51] |
| 7KJ4 | 1069 | 0.4 | 3.4 | Three | (Xiao et al., 2021) [51] |
| 7KMS | 1086 | 0.0 | 3.6 | Three | (Zhou et al., 2020) [52] |
| 7KMZ | 1085 | 0.0 | 3.6 | Two | (Zhou et al., 2020) [52] |
| 7KNB | 1083 | 0.0 | 3.9 | One | (Zhou et al., 2020) [52] |
| 7KNE | 1083 | 0.0 | 3.9 | One | (Zhou et al., 2020) [52] |
| 7KNH | 1085 | 0.0 | 3.7 | Two | (Zhou et al., 2020) [52] |
| 7KNI | 1082 | 0.0 | 3.9 | Three | (Zhou et al., 2020) [52] |

### 2.2 Computing the change in ACE2 binding affinity due to mutation

We used the BeatMusic method (default setting) [53], which has shown very promising accuracy in independent benchmarks [54,55], to compute the binding affinity (ΔΔG_bind_, in units of kcal/mol) for the 21 selected structures. For each input structure, we define the two interacting partners – one being the three chains of S-protein and the other being ACE2. The method uses coarse-grained models to compute protein-protein interactions via statistical potentials. If more than one chain of identical sequence is present in one partner, the method introduces mutations in all of them, and thus the ΔΔG_bind_ represents the total change in binding free energy due to mutation, with a negative value representing an increase in binding affinity. The relative solvent accessibilities (RSA) of the mutated residue sites were also calculated using all 21 structures as input. BeatMusic computes RSA of the mutated sites in the complex and apo states as the ratio of the solvent accessible surface in the selected structure (using DSSP) [56] and in a corresponding extended tripeptide Gly-X-Gly [53,57]. Previous analysis has shown that the solvent accessibility of individual sites (likely to affect estimates of binding affinity) such as D614 varies substantially in different experimental structures, reflecting the conformational dynamics discussed above [21], which suggests that local site heterogeneity may affect mutation estimates.

### 2.3 Mutation group comparisons

Mutations in proteins are on average more likely to be destabilizing due to the optimization for a fold stability [58,59]. Both the apo-protein and the complex are likely to experience loss of stability upon mutation, but the net impact on binding affinity depends on the relative stability change of the complex and apo-protein. Computational methods to estimate mutation effects suffer from a number of biases relating to the extrapolation from to a specific static wild type structure to a (usually unknown) mutant structure (missing information and structural heterogeneity), the accuracy of the model’s physics, and the biases in the datasets used to train them, and they are neither very accurate nor precise (results depend on method and structure used) for individual mutations [32–34,60]. We have previously demonstrated how both systematic and random errors in such methods can be reduced by comparing the averages of case and control mutation groups, e.g., the typical properties of pathogenic mutations relative to the background of similar but random mutations but have not applied this protocol to SARS-CoV-2 mutations affecting S-protein ACE2 binding where it would be equally relevant [61,62].

To do so, we compared the average and standard deviations ΔΔG_bind_ of groups of reported natural mutations against various other groups of mutations. We computed and analyzed ΔΔG_bind_ for all possible N x 19 mutations (~18,000 to 21,000 mutations per structure) of all 21 structures, for a total of approximately 400,000 ΔΔG_bind_ data points.

We then divided mutations into seven different groups (**Table S1**): (*a*) all possible mutations in the full S-protein (N x 19); (*b*) all possible mutations in sites at the S-protein-ACE2 binding interface, defined as having a BeatMusic-computed decrease in solvent accessibility in the complex of at least 5%; (*c*) all naturally observed mutations in the alpha, beta, gamma, delta, omicron, lambda, mu, kappa, iota, eta, zeta, theta, epsilon variants, and other consistently identified circulating mutations compiled from UniProt ID P0DTC2 as of 15 April 2022 (**Figure 1**, giving 79 substitutions in 71 different sites, of which 70 mutations could be computed due to available site coordinates in at least one structure); (*d*) all possible substitutions at sites known to have experienced natural mutation (i.e., evolvable sites); (*e*) mutations observed in the most prevalent variants indicative of combined high fitness (alpha, beta, gamma, delta, and omicron); (*f*) other mutations observed but not typical of the most prevalent variants; and (*g*) mutations present in the omicron variant.

**Figure 1.**
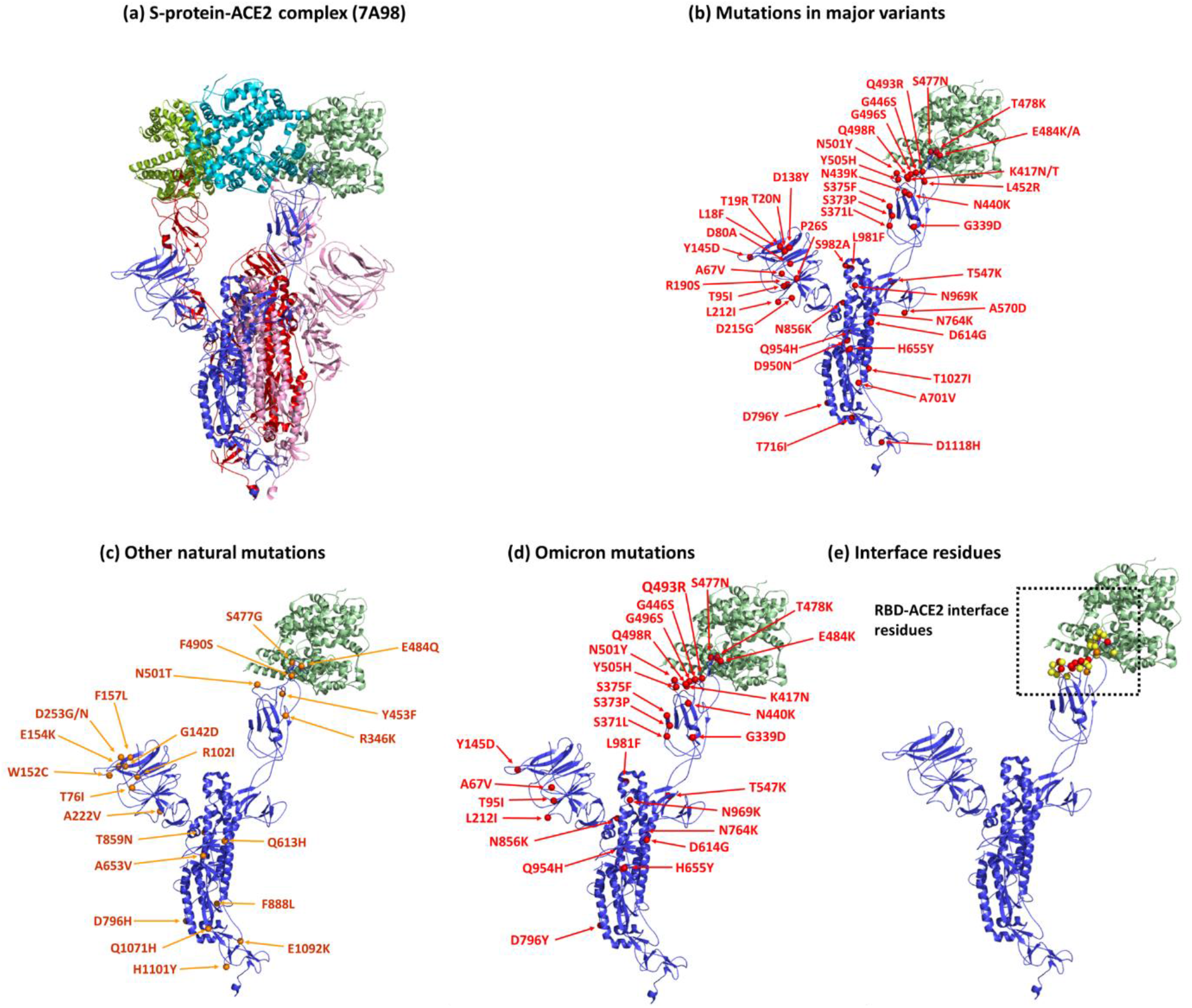
Structure and mutations of the SARS-CoV-2 S-protein. **(a)** The S-protein trimer with its three chains (blue, red, and purple) and three ACE2 proteins (dark green, cyan, and light green). **(b)** Mutations seen in major SARS-CoV-2 variants (alpha, beta, gamma, delta, omicron, lambda, mu, kappa, iota, eta, zeta, theta, epsilon). **(c)** Selected additional mutations repeatedly seen in SARS-CoV-2 sequences. **(d)** Mutations seen in the original BA.1 omicron variant. **€** S-protein-ACE2 interface residues (yellow), with residues in red and orange indicating mutation sites observed in major variants and in other variants, respectively.

The group of all natural mutations include the mutations seen in major variants, mutations of interest, mutations under monitoring and other common naturally occurring mutations. Of course, this selection is far from exhaustive as very many single substitutions have been seen in sequences across the world, but they have uncertain functional impacts and have not been fixated in the broader populations, and thus would not be of special interest to our investigation. The sets *e* and *f* are subsets of set *c* i.e., sets “*e* + *f* = set *c*”, whereas set *g* is a subset of set *e*. In other words, set f = *c* – *e*, i.e., sets *e* and *f* are disjoint. These data were compiled from WHO [63], UniProt [64], and the CDC [65] web pages of mutation monitoring.

### 2.4 Sensitivity to binding stoichiometry

The studied complexes represent S-proteins bound to three, two or one ACE2 protein, binding at 3, 2, or 1 of the RBDs of the trimer S-protein. While the physiological stoichiometry of S-protein-ACE2 complexes may potentially vary (1:1 being probably more likely, given the many other constituent proteins in the human cell membrane) inclusion of these structures enables us to test the sensitivity of our results to such changes in binding stoichiometry.

To this end, we performed computations using three different PDB structures 7DX6, 7DX8 and 7KJ4 where the S-protein is bound to 1, 2 and 3 ACE2 respectively (**Figure S1**). We found that only a few protein sites are affected: Correlation coefficients of linear regression (R) was 0.99 for 1 versus 2 ACE2 proteins, 0.91 for 1 versus 3 ACE2 proteins, and 0.91 for 2 versus 3 ACE2 proteins (**Figure S1**), i.e., the vast majority of mutation effects are not impacted by ACE2 stoichiometry. Mutations lying far from the central regression line in **Figure S1** are few and mostly at the interface (**Table S2**) for R=0.99, whereas in case of R=0.91, there are more sites affected and most of these are not at the interface, suggesting that the mutation effect is generally not impacted by ACE2 stoichiometry.

### 2.5 Experimental ACE2 binding data

The computed ΔΔG_bind_ values were compared with the experimentally reported ACE2 binding affinities for mutations in the RBD by Bloom’s group [30]. These data points that were not reproduced, defined as residuals > 1, or where data for one replicate was missing, were excluded, to ensure that only reproducible experimental data points were used. We additionally removed data points with binding effects in either replicate more negative than −4.5 to avoid values near the detection limit, as inspection of the data shows artificial clustering of data points in this region which will affect an analysis erroneously. These curations also led to exclusion of stop codon mutations and also the mutation sites that were not present in any of the 21 studied structures were excluded, in total removed 831 (3390 data points left) of the original 4221 experimental data points from consideration.

### 2.6 Statistical analysis

For the statistical analysis to determine the significance of the binding effects, we compared the results for the average ΔΔGbind and standard deviations of the defined mutation groups, and the RSA for each group of mutations using student’s t-tests for same mean. The average ΔΔG_bind_ was calculated in the following ways: (i) for each of the seven mutation groups in each structure, (ii) for the same natural mutation in 21 structures, (iii) for the average of 21 averages of the 21 structures for each of the seven mutation groups, (iv) for the same mutation in RBD for 21 structures for comparison with experimental binding and expression data. This protocol reduces the impact of both random and systematic errors and removes some of the uncertainty of single-mutation estimates based on a single structure, as discussed further below.

## 3. Results and discussion

### 3.1 Computed binding affinities correlate with the experimental data

To understand the overall trends in the computed numbers, we compared our binding affinities to the recent experimental binding data specifically for RBD mutations [30]. The experimental values were multiplied by −1 to make them uniform with the computed ΔΔG_bind_ values so that negative values indicate increased binding affinity. For comparison, the RBD mutations were categorized into full-length RBD, mutations at the RBD-ACE2 interface, natural mutations within the RBD, and natural mutations at the RBD-ACE2 interface.

Despite the expected substantial variations for individual mutations, the predicted and experimental values show similar trends (**Figure 2a, 2b**). The random RBD and interface mutations showed more decreased ACE2 binding than the natural mutations in the RBD and at the interface. The average data (**Figure 2b**) showed clear separation between random RDB and natural mutations, and no clear difference between natural mutations at the interface and within the RBD. The similar trends of computation and experiment suggest that BeatMusic can predict the general experimental tendencies of the groups, even when results are unreliable for single mutations; this explains the need for mutation group comparisons.

**Figure 2.**
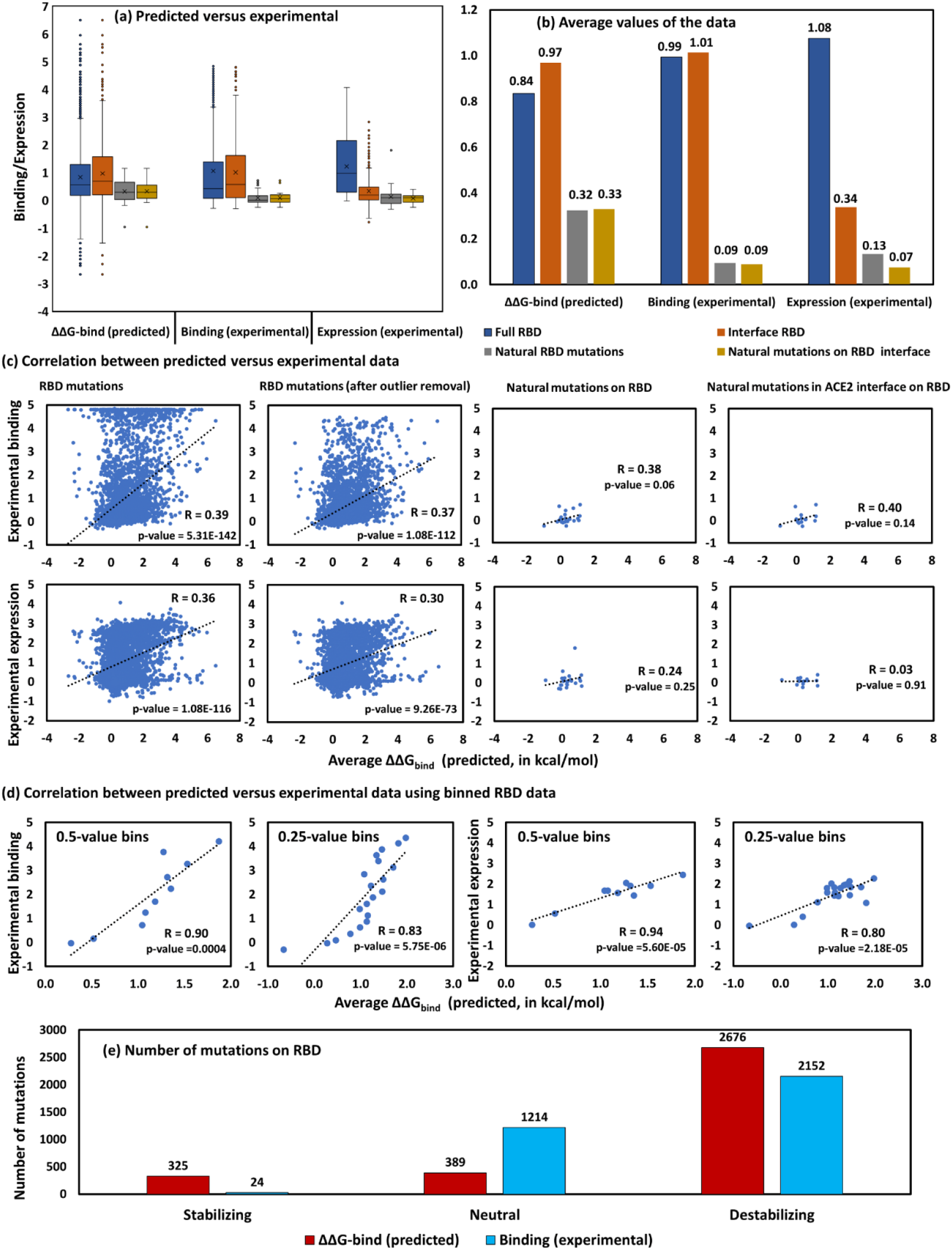
Correlation of computed average ΔΔG_bind_ of RBD mutations with experimental binding and expression. **(a)** Comparison of computed versus experimental data. For computed ΔΔG_bind_, the average for the 21 structures was used. **(b)** Average of the computed and experimental data for each mutation group. For the analysis in panels **(a)** and **(b)**, values which were not experimentally available were removed (3797 data points). **(c)** Correlation between computed average ΔΔG_bind_ and experimental data (3797 points in the left panel and after outlier removal 3390 points). **(d)** Correlation between computed and experimental data using 0.5- and 0.25-value bins (3390 data points). **(e)** Number of mutations on RBD belonging to stabilizing, neutral and destabilizing binding groups for computed and experimental data.

Scatter plots of the computed ΔΔG_bind_ and experimental binding and expression data [30] are shown in **Figure 2c**. We computed the average ΔΔG_bind_ for each RBD mutation from the 21 PDB structures and used these average ΔΔG_bind_ values. We found significant correlation between computed average ΔΔG_bind_ values and experimental binding (R = 0.39) and expression data (R = 0.36). Removal of outliers did not increase correlation substantially. Correlation was also seen specifically for natural mutations present on RBD (R = 0.38 for binding and R = 0.24 for expression, **Table S3**). Though the R-values for the natural mutations are not significant, they are significant for the full RBD mutations, with the same direction in all analyses, indicating that the underlying physics of the effects is captured but that uncertainties in individual mutations and structural variations make the results imprecise. Accordingly, computations can reproduce reasonably the trend if accounting for heterogeneity in the structural data. We note that the experimental expression and ACE2 binding also correlate, which may be due to a common causal property of the mutations, or S-protein concentration impacting the observed ACE2 binding. However, the R-values are not very large despite significant p-values, as further analyzed below.

The experimental values analyzed above are highly skewed (**Figure 2c**). To undo this effect, we grouped the data into bins of 0.5- and 0.25-width for the experimental binding and expression data, calculated average binding and expression data for each such bin, and plotted correlation with the computed ΔΔG_bind_ averaged from bins as described in the methods section. As shown in **Figure 2d**, this produces highly significant correlations between both binding and expression and computed structure-averaged ΔΔG_bind_. The R-values for experimental versus predicted binding effects are 0.90 and 0.83 for 0.5- and 0.25-value bins, respectively, which is surprisingly accurate but also reflects a massive compression of the data. The R values for expression versus predicted ΔΔG_bind_ are 0.94 and 0.80 for 0.5- and 0.25-value bins. Removal of the overrepresented data via bins substantially improved the correlation (**Figure 2c**).

We also analyzed the RBD mutation data for the impact of mutations on ΔΔGbind ranked by magnitude to visualize the broadness, i.e., to understand whether the effect in a given variant is most likely dominated by a few or broadly many mutations (with the limitation that epistasis is not accounted for). We classified the data into improved binding (< −0.1), neutral (−0.1 to 0.1) and destabilizing (>0.1) mutations and compared with the experimental binding data by keeping the same quantitative divisions (**Table S4** and **Figure 2e**). Importantly, and further validating the performance of the protocol, the predicted and experimental binding values follow a similar pattern, with most mutations showing negative or nearly neutral effects, and a very small number of mutations having beneficial effects on ACE2 binding.

### 3.2 Computed ACE2 binding: Comparing natural and random mutations

The results of the ΔΔG_bind_ analysis for the seven groups of mutations for all 21 S-protein-ACE2 structures are shown in **Figure 3**. The results for all possible mutations in the protein (blue boxes to the left) represent completely random mutations, including mutations not likely to occur in the wild due to various selection pressures and in sites far from the RBD. Most mutations have positive values, suggesting decreased binding affinity for all seven mutation groups in all 21 structures. This may partly be due to a bias in the data set used to train the method towards complexes evolved or designed towards high affinity, or it can be a real effect indicating that the S-protein itself is optimized towards ACE2 binding, such that most mutations tend to impair this optimized affinity. The experimental data by Starr et al.[30] are consistent with such an average tendency indicating that the effect is not simply due to a method bias, but the systematic errors inherent to any computational method render the direct values not very meaningful.

**Figure 3.**
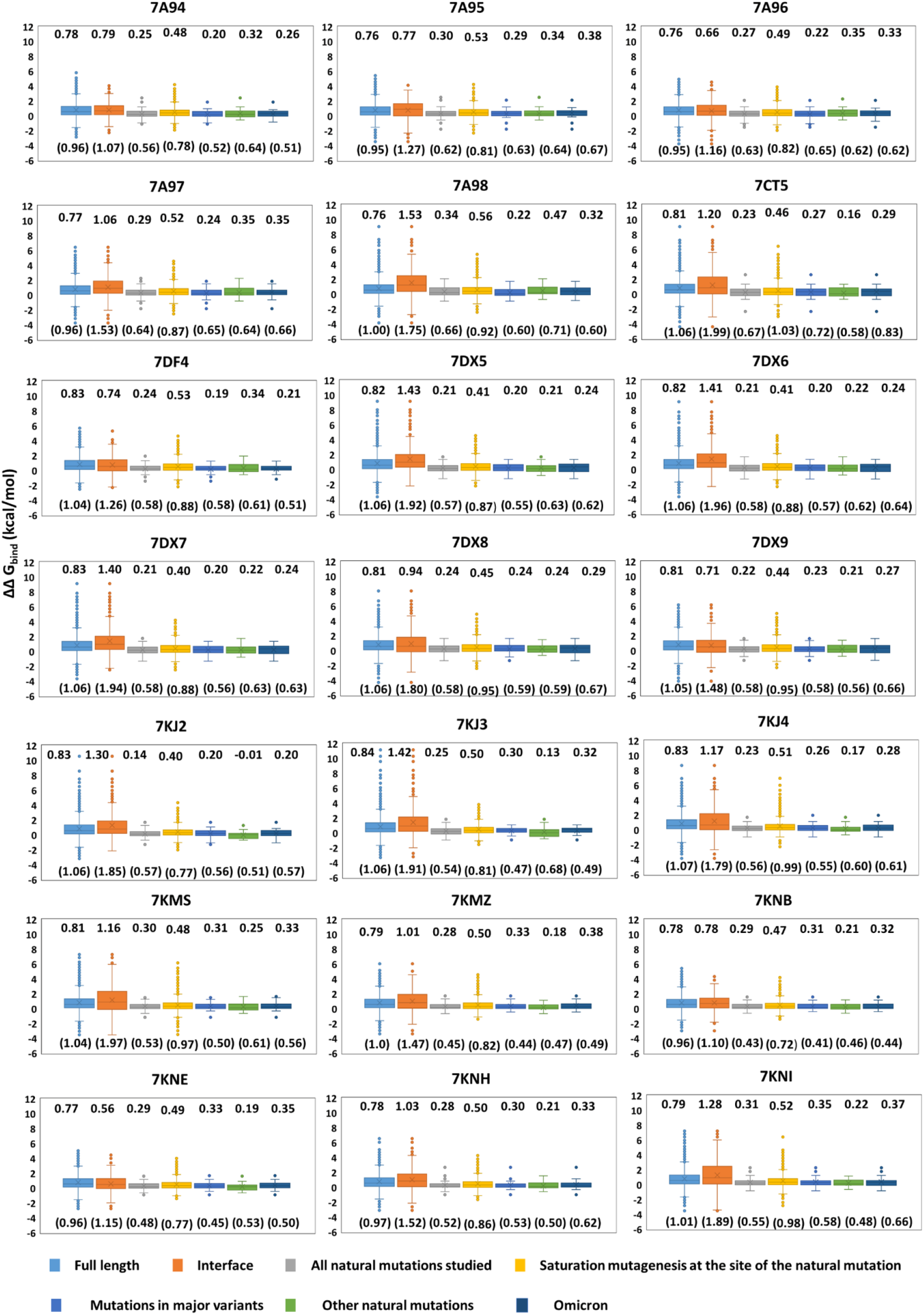
Change in binding free energy (ΔΔG_bind_) upon mutation for the 21 S-protein-ACE2 complexes. The average ΔΔG_bind_ for each mutation group is indicated in the top of each panel, and standard deviations are shown in brackets in the bottom of each panel.

Instead, comparison of the averages of the different groups removes systematic errors in the method. Using this approach, the natural mutations, regardless of group, showed consistently better ACE2 binding than random full-protein mutations (light blue boxes to the left) and mutations at the interface (orange). More interestingly, all possible mutations at sites of known natural mutations (**Figure 3**, yellow) give values intermediate between actual natural mutations and all S-protein mutations, i.e., the fixated natural mutations are predicted to bind ACE2 better on average than other mutations even in the same sites, located similarly relative to ACE2.

We found no significant difference between the mutations for the major variants and omicron, indicating that the variant mutations combine to relatively similar ACE2 binding compared to random mutations in the evolvable and exposed sites. While these estimates represent the pure individual mutation contributions which may be modulated somewhat by epistasis (interactions between sites that change the combined effect on binding) the tendency is supported by estimates of ACE2 binding of delta and omicron being relatively similar [29], although others found a somewhat smaller binding affinity for omicron than delta [28]. The corresponding analysis for pairs of mutation groups in **Figure S2** for all 21 structures shows very good agreement for the main batch comparisons, despite large variations in some individual mutations, illustrating the importance of considering groups of mutations (to increase precision) rather than uncertain single-mutation estimates in computational studies.

### 3.3 Taking into account structural heterogeneity

**Figure 4a** shows the ΔΔG_bind_ values for the 70 natural mutations in the S-protein averaged for all structures where the site coordinates are present. Whereas the direct effect values are not significant due to systematic model errors, the difference in effect between mutations have reduced systematic error but are still not precise (**Figure 4a**). However, they illustrate how large differences in individual effects average out to small group differences in **Figure 3**. Thus, the combination of structure averaging and group comparisons considerably strengthens any such computational protocol. If we consider 0±0.5 kcal/mol as the cut-off for neutral mutations, most of the mutations fall in that range. Outside the ACE2 interface region we estimate large negative effects of Y145D and W152C. Since these sites do not form a part of RBD motif, they do not directly interact with ACE2, yet the effect is predicted repeatedly by different structures. N501Y gives consistently stronger binding, relative to other prominent mutations, consistent with experimental kinetic and affinity data [24].

**Figure 4.**
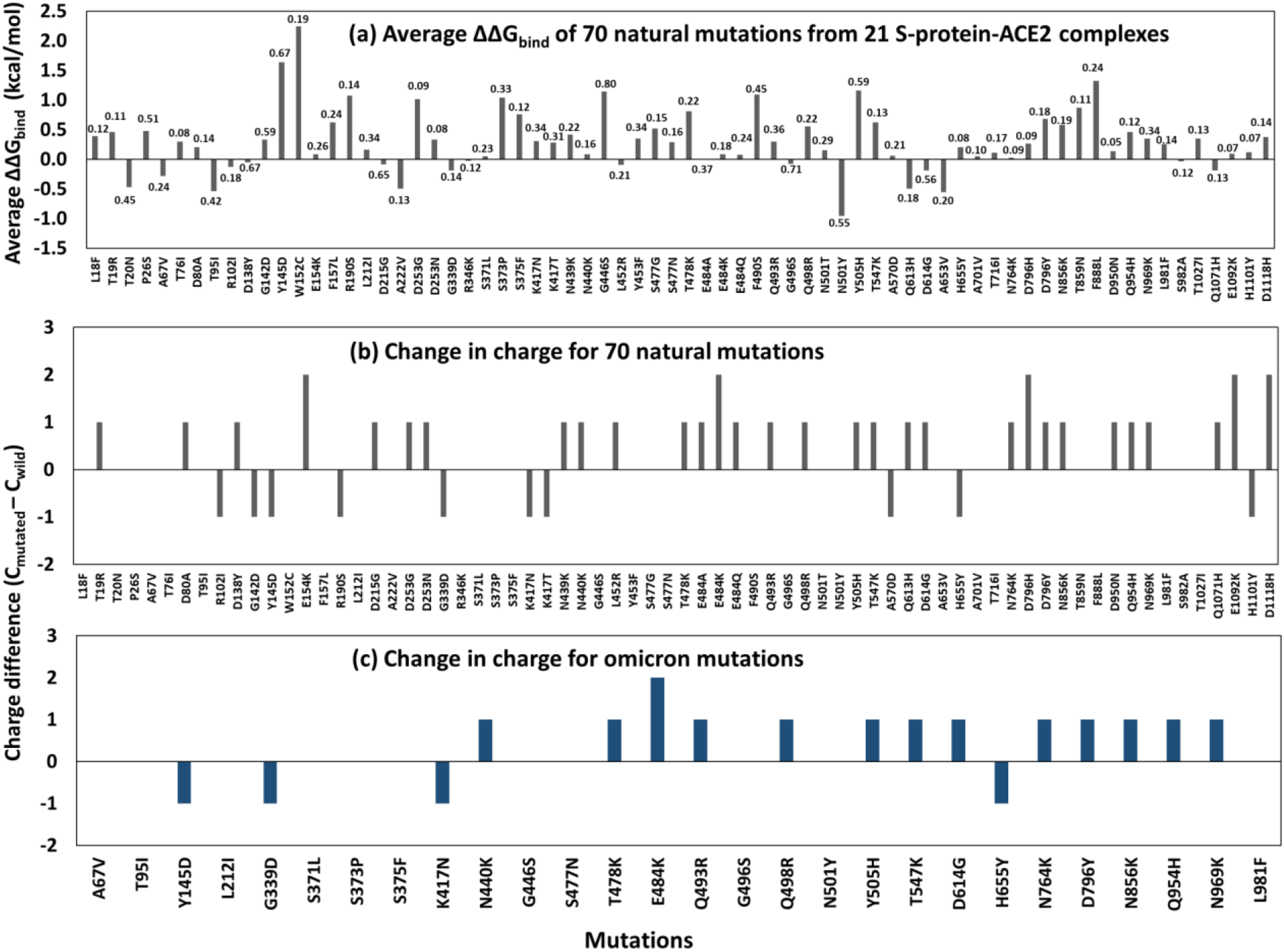
Average ΔΔG_bind_ and change in charge values for natural mutations. **(a)** Average ΔΔG_bind_ calculated from 21 structures for the natural mutations with standard deviations written for each bar. **(b)** Change in charge (C_mutated_ – C_wild_) for the 70 studied natural mutations. **(c)** Change in charge for omicron mutations.

To illustrate the impact of structure choice, **Figure S3** shows the variation in ΔΔG_bind_ values for some selected natural mutations for all studied structures. For example, D614G has positive or negative effect on ACE2 binding almost equally distributed depending on the chosen structure, whereas some mutation effects are insensitive to structure use. This result originates in the large conformational variability of some surface sites, with D614 being a notable example [21]. For this reason, computational studies based on a single structure may give misleading results, as the structure’s site conformation may not be representative of the full knowledge we have from cryo-EM structures of the S-protein.

We also note that the change in charge of the natural mutations (**Figure 4b**) including omicron (**Figure 4c**), shows that many mutations lead to increased charge. Out of 70 natural mutations, 30 cause increased charge, whereas only 10 reduce charge and 30 are charge neutral. In omicron, out of 28 mutations (present in structures analyzed), 13 increase the charge and 4 decrease the charge. This could suggest that the mutations that cause positive charge tend to be more often associated with stronger binding to the negatively charged surface of ACE2 [21]. Earlier studies have reported that charge increment can enhance ACE2 binding affinity [24,26,27,66–69]. The charge-increasing mutations N439K, N440K, E484A, E484K, E484Q, Q493R and Q498R show near neutral effect (with 0.5 kcal/mol cut-off for neutral, **Figure 4a**), but considering the systematic tendency towards weaker binding, this mainly indicates a stronger binding relative to random mutations as can be seen in group comparison in **Figure 3**.

As noted above, mutation effects sometime depend substantially on the structure used as input (**Figure S3**). To account for this structural heterogeneity, we can average data for the mutations across all the identified structures fulfilling the inclusion criteria. The resulting mutation effects represents the effect in a combined conformational space of the cryo-EM structures, which is missing when using a single structure. The conformations of the S-protein can be very dependent on conditions such as pH [52], which supports this type of analysis.

A corresponding total summary of structure-averaged mutation effects is shown in **Figure 5a**, with **Figure 5b** showing the statistics (t-test for same mean) of group pair-comparisons. The most important significant differences are, as discussed above, all the natural mutation groups (grey, blue, green, dark blue) vs. all possible S-protein mutations (light blue; left box), spike-ACE2 interface (orange) and all possible mutations in evolvable sites (yellow) (**Figure 5a**), with comparisons between groups of natural mutations being insignificant (except omicron). The S-protein-ACE2 interface region is the contact point for ACE2 and therefore any random disturbance in this region is expected to have large effect on ACE2 binding as observed in the orange bar in **Figure 5a**. The omicron mutations as a group show significantly decreased ACE2 binding compared to other natural mutations, including those in major variants, but the effect is small and could thus explain the discrepancy between experimental findings, with studies finding either that omicron has less [28] or similar [29] binding to ACE2 compared to other variants. A recent computational study indicated that the RBD-ACE2 binding increases somewhat in the omicron variant compared to wild-type RBD [70], which is not seen in our comparison but the effects are small and the methods quite different, so this is not so surprising.

**Figure 5.**
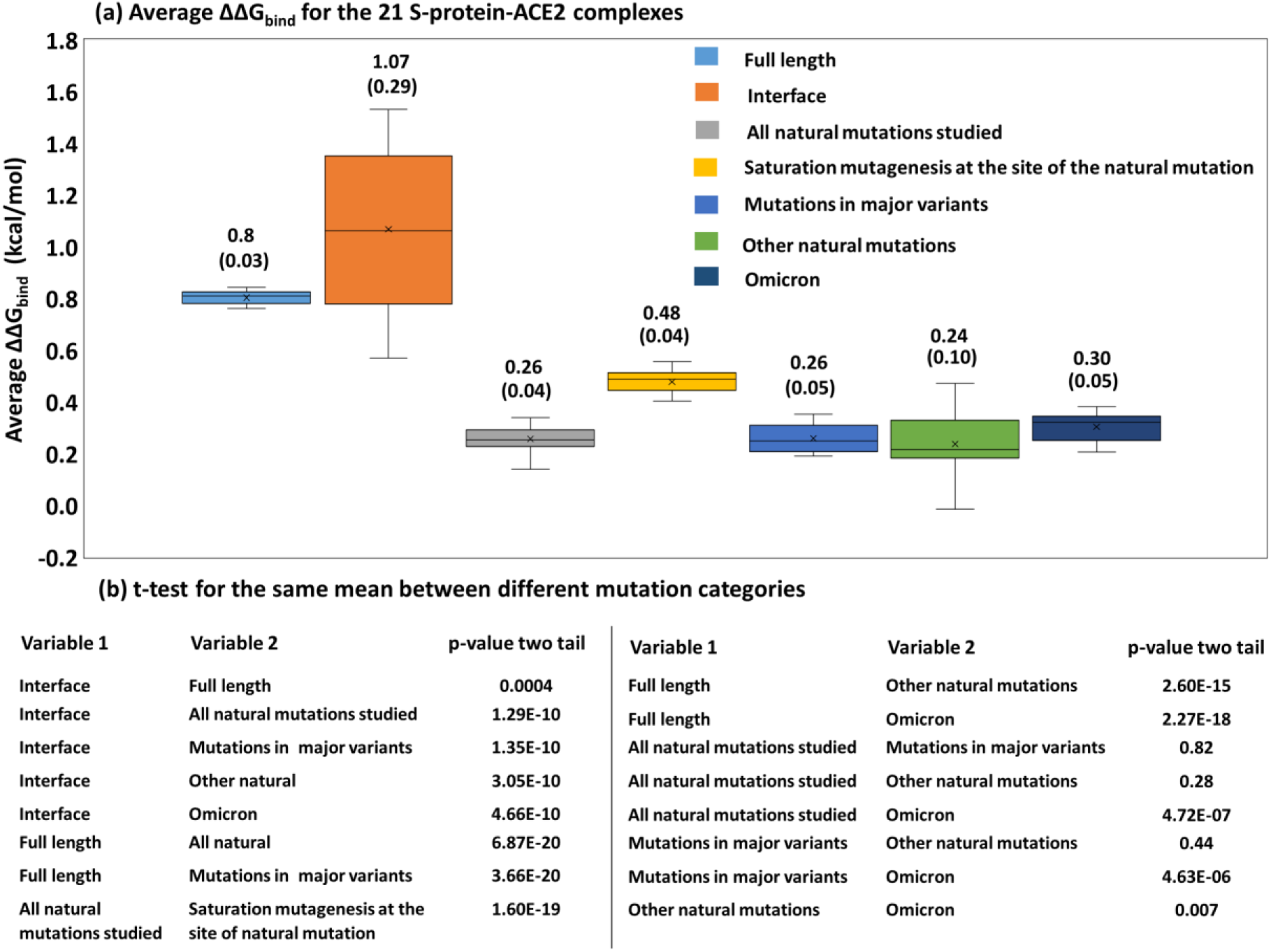
Comparison of the average ΔΔG_bind_ values. **(a)** ΔΔG_bind_ values averaged over the 21 S-protein-ACE2 complexes for seven groups of mutations, with standard deviations in brackets. **(b)** p-values of group comparisons using t-test for same mean for the data in panel **(a)**.

### 3.4 Solvent accessibility of spike-ACE2 complexes

An analysis of the RSA for the 21 PDB structures is shown for the complex in **Figure 6** and after removing ACE2 in **Figure S4**. As seen in **Figure 6**, the average RSA of S-protein-ACE2 complexes of natural mutations is consistently high in comparison to all possible mutations for all the 21 structures. This trend was also noted for natural mutations relative to the mutations at the ACE2 interface (except 7A96). The natural mutation sites and the S-protein-ACE2 interface are thus relatively solvent exposed, even in the ACE2 complex state, compared to random sites. We note however that some of the natural mutations exhibited very varied solvent accessibility, due to variations in the cryo-EM structures, which again documents the need for using aggregate “ensemble” results from multiple structures.

**Figure 6.**
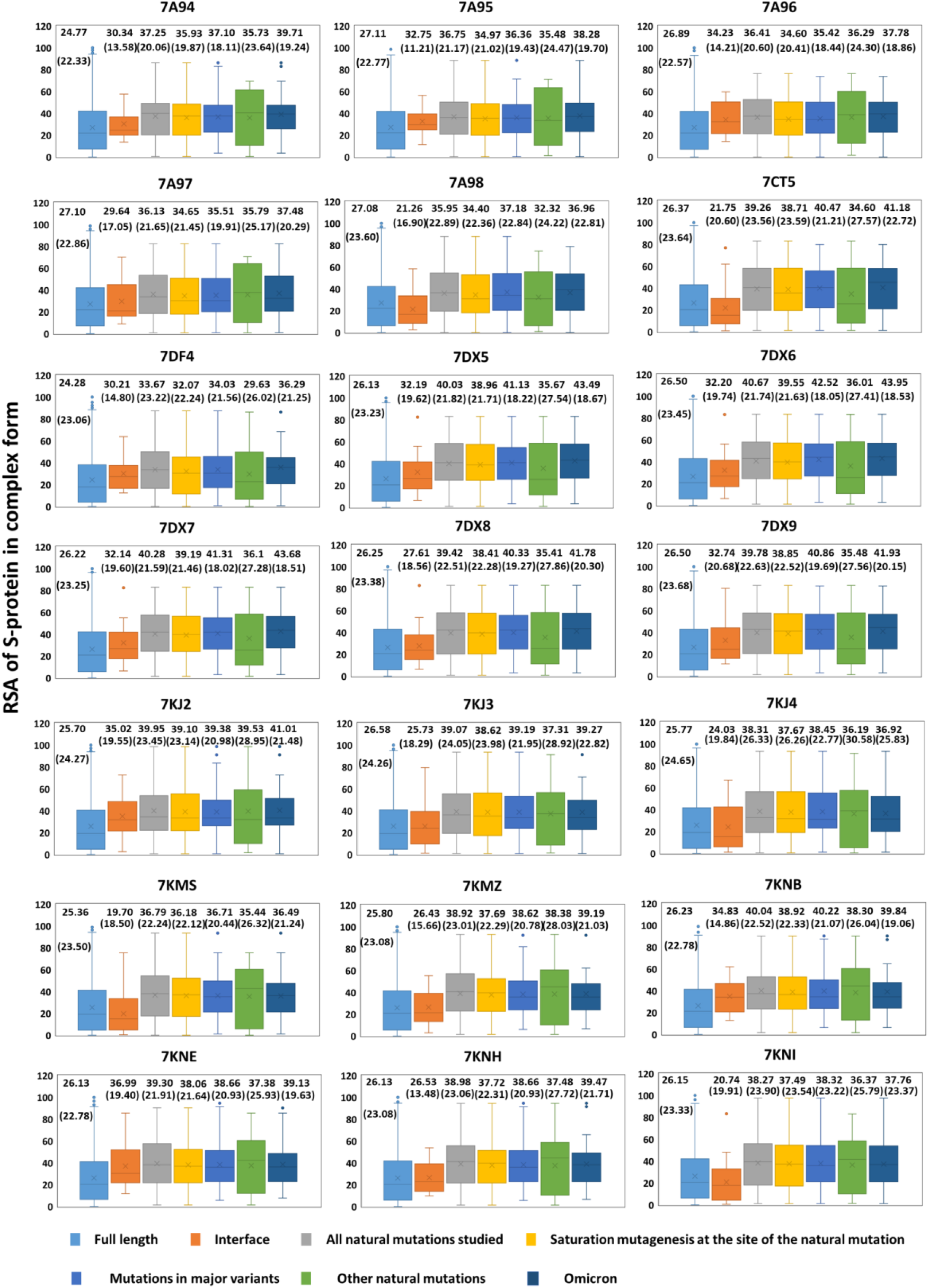
Residue solvent accessibility for the 21 S-protein-ACE2 complexes. The average RSA and standard deviation for each mutation group are indicated in the plots (standard deviation represented inside bracket). The saturation mutagenesis for the site of natural mutations (yellow) is the data averaged over 19 possible mutations for the same site and therefore differ slightly from the group of all natural mutations (grey).

The average RSA of 21 S-protein structures after removing ACE2 showed similar behavior of consistently high solvent exposure for natural mutation comparative to all possible mutations (**Figure S4**). The interface residues show a clear increase in solvent accessibility for all 21 structures, as anticipated, on average of ~10 to 23, when ACE2 was removed. Solvent exposure is important for the mutation tendency, but this is usually due to these sites being more functionally neutral, thus evolving faster than structurally constrained buried sites [71], whereas in the S-protein it is due to many solvent-exposed sites being subject to positive selection, which also increases evolution rates.

We note that mutations that favor binding to one protein (e.g., any antibody) are likely to more often also favor binding to another protein (e.g., ACE2), such that classes of mutations are likely to bind better to both, giving a correlation between ACE2 and antibody binding. Our results suggest that ACE2 binding has been almost maintained or slightly relaxed by a combination of many mutations having different magnitudes of their effects in a plausible trade-off with antibody evasion. However antibodies bind in very different parts of the protein [72] and thus any use of a single antibody as representative for such correlations should be taken with substantial caution. A recent study identifies 377 human monoclonal antibodies for S-protein with ~80 of them binding to RBD, indicating that the antibody sites are very diverse [10]. A study on epistasis [40] relied on one antibody as representative of this entire epistatic relationship with ACE2 and also only one experimental structure for computing the results. We consider this unlikely to be a strong estimate of whether epistasis is important or not and invite more detailed studies into epistasis, including both intra-S-protein and inter-gene epistasis at both the RNA and amino acid level, preferably with group comparisons and averaging over multiple equally valid cryo-EM structures to reduce noise, as proposed in the present study.

## 4. Conclusions

Computational structure-based screening of new S-protein mutations for ACE2 binding is required to rationalize virus function directly from protein structure and may eventually aid early detection of potentially concerning variants. However, substantial systematic and random errors in the models and variations in the cryo-EM structures used as input lead to issues with both accuracy and precision. In this work, we investigated how computations may be used to estimate ACE2 binding effects of new SARS-CoV-2 mutations, using mutation group comparisons to reduce systematic errors and the richness of cryo-electron microscopy structures of the S-protein to our advantage as an ensemble to reduce noise in individual mutation estimates. The protocol gives reasonable trend agreement with experimental mutation data for the RBD, but is then computationally extended to the full mutation landscape of 21 different protein structures. Accordingly, the main findings of this work include:

- The computed mutation effects are often highly sensitive to the structure used as input, with e.g., the effect of D614G on ACE2 binding being entirely dependent on structure used.
- We propose a protocol that averages out the structural heterogeneity to give more robust estimates, and show how comparisons of differences in average effects can produce significant trends in the ACE2 binding behavior for groups of mutations.
- We find that the S-protein-ACE2 interface region is sensitive to mutation effects, but more importantly that the natural mutations have better binding than random mutations in evolvable sites and the full S-protein, quantifying the presence of an optimization of ACE2 binding but also providing point estimates for mutations not studied experimentally.
- We found that the variant mutations as groups summed to relatively similar binding effects, despite substantial individual variations to both sides. This suggests that the protocol is reasonably accurate in estimating combined mutation effects despite the absence of epistasis in the model.
- Omicron mutations exhibited slightly lower binding affinity as compared to mutations in other major variants, which is consistent with experimental studies finding somewhat less [28] or similar [29] binding to ACE2 compared to delta.
- Many natural mutations have led to positive charge increase, which plausibly contributes to strong electrostatic interaction with the negatively charged surface of ACE2.

Our protocol and structure-averaged point estimates may be of use in future models that estimate the S-protein-ACE2 binding effects of new mutations and rationalize evolutionary trade-offs of individual mutations with regards to antigenic drift, which we believe is more difficult to describe due to the complexity and antibody-S-protein interactions. In addition to outlining a substantially better computational protocol to estimate ACE2 binding affinity, the detailed genotype phenotype map provided for the full mutation space averaged over 21 experimental cryo-EM structures is the main outcome of our study, which we expect to use for new models with improved predictability of virus evolution.

## Acknowledgements

RM, ST and RKV acknowledge IIT Bhilai for the supporting this work via Research Initiation Grant (RIG), project code 2005900. ST acknowledges Ministry of Education, Government of India for research fellowship.

## Data availability

The Supporting Information file contains additional analysis of the binding data.

