## Supplementary file for "Modelling SARS-CoV-2 spike-protein mutation effects on ACE2 binding"

*Rukmankesh Mehra: 0000-0001-6010-1514*

*Kasper P. Kepp: 0000-0002-6754-7348*

**Table S1. Categories of mutations analyzed in the present study.** Data were compiled as of April 15, 2022, from the UniProt ID P0DTC2 and CDC databases. All mutations studied are substitutions (not e.g., deletions). Set *a* is the set of all possible mutations in the full S-protein. Set *b* is the set of known mutations at the S-protein-ACE2 interface as defined in the main text. Set *c* ("all natural mutations studied") consists of all substitutions of the alpha, beta, gamma, delta, omicron, lambda, mu, kappa, iota, eta, zeta, theta, and epsilon variants and additional mutations commonly identified in circulating sequences (79 substitutions in 71 different sites). 70 of these substitutions could be modelled from at least one of the 21 PDB structures (some mutations L5F, S13I, G75V, Q677H, N679K, P681H/R, V1176F, and G1219V occur in sites that are missing in the structures). Set *d* consists of all mutations possible in the sites of the studied natural mutations in set *c*. Set *e* ("mutations in major variants") contains the natural mutations from the alpha, beta, gamma, delta and omicron variants (51 substitutions in 48 different sites). Set *f* ("Other natural mutations") consists of 28 substitutions in 27 sites and is a subset of the natural mutations set *c* not characteristic of the major variants. Set *g* represents the substitutions in the omicron variants, a subset of set *c*, taken from CDC database (30 amino acid substitutions in 30 different sites).

| **Set** | **Mutation categories** | **Mutations** |
|---|---|---|
| *a* | S-protein | Saturation mutagenesis |
| *b* | Spike-ACE2 interface (29 in total) | R403, D405, K417, V445, G446, Y449, Y453, L455, F456, Y473, A475, G476, S477, N481, E484, G485, F486, N487, Y489, F490, Q493, S494, G496, Q498, T500, N501, G502, V503, Y505 |
| *c* | All natural mutations studied (79 in total, in 71 different sites) | L5F, S13I, L18F, T19R, T20N, P26S, A67V, G75V, T76I, D80A, T95I, R102I, D138Y, G142D, Y145D, W152C, E154K, F157L, R190S, L212I, D215G, A222V, D253G/N, G339D, R346K, S371L, S373P, S375F, K417N/T, N439K, N440K, G446S, L452R, Y453F, S477G/N, T478K, E484K/A/Q, F490S, Q493R, G496S, Q498R, N501Y/T, Y505H, T547K, A570D, Q613H, D614G, A653V, H655Y, Q677H, N679K, P681H/R, A701V, T716I, N764K, D796H/Y, N856K, T859N, F888L, D950N, Q954H, N969K, L981F, S982A, T1027I, Q1071H, E1092K, H1101Y, D1118H, V1176F, G1219V |
| *d* | Saturation mutagenesis at the site of natural mutations | All possible mutations at the natural mutation sites in (c) |

| | | |
|---|---|---|
| *e* | Mutations in major variants (51 in total, in 48 different sites) | L18F, T19R, T20N, P26S, A67V, D80A, T95I, D138Y, Y145D, R190S, L212I, D215G, G339D, S371L, S373P, S375F, K417N/T, N439K, N440K, G446S, L452R, S477N, T478K, E484K/A, Q493R, G496S, Q498R, N501Y, Y505H, T547K, A570D, D614G, H655Y, N679K, P681H/R, A701V, T716I, N764K, D796Y, N856K, D950N, Q954H, N969K, L981F, S982A, T1027I, D1118H, V1176F |
| *f* | Other natural mutations (28 in total, in 27 different sites) | L5F, S13I, G75V, T76I, R102I, G142D, W152C, E154K, F157L, A222V, D253G/N, R346K, Y453F, S477G, E484Q, F490S, N501T, Q613H, A653V, Q677H, D796H, T859N, F888L, Q1071H, E1092K, H1101Y, G1219V |
| *g* | Omicron (30 substitutions in 30 different sites) | A67V, T95I, Y145D, L212I, G339D, S371L, S373P, S375F, K417N, N440K, G446S, S477N, T478K, E484K, Q493R, G496S, Q498R, N501Y, Y505H, T547K, D614G, H655Y, N679K, P681H, N764K, D796Y, N856K, Q954H, N969K, L981F |

**Table S2. Outlier analysis.** Analysis of the residues lying away from the regression lines in correlation plots of **Figure S1** to understand whether they are present at the S-protein-ACE2 interface (indicated by Yes or No).

| **1 ACE2 versus 2 ACE2 complexes (7DX6 versus 7DX8; R=0.99)** | | | **1 ACE2 versus 3 ACE2 (7DX6 versus 7KJ4; R=0.91)** | | | **2 ACE2 versus 3 ACE2 (7DX8 versus 7KJ4; R=0.91)** | |
|---|---|---|---|---|---|---|---|
| | | | 27 | No | | 131 | No |
| 528 | No | | 89 | No | | 133 | No |
| 489 | Yes | | 133 | No | | 135 | No |
| 493 | Yes | | 135 | No | | 139 | No |
| 500 | Yes | | 139 | No | | 200 | No |
| 502 | Yes | | 200 | No | | 451 | No |
| 505 | Yes | | 453 | No | | 453 | No |
| | | | 483 | No | | 483 | No |
| | | | 495 | No | | 495 | No |
| | | | 497 | No | | 515 | No |
| | | | 515 | No | | 524 | No |
| | | | 527 | No | | 601 | No |
| | | | 601 | No | | 720 | No |
| | | | 882 | No | | 891 | No |
| | | | 476 | Yes | | 908 | No |
| | | | 486 | Yes | | 1082 | No |
| | | | 489 | Yes | | 476 | Yes |
| | | | 493 | Yes | | 486 | Yes |
| | | | 496 | Yes | | 489 | Yes |
| | | | 498 | Yes | | 496 | Yes |
| | | | 500 | Yes | | 498 | Yes |
| | | | 502 | Yes | | 500 | Yes |
| | | | 505 | Yes | | 502 | Yes |

**Table S3. Comparison of experimental (Bloom data) versus predicted data for the natural mutations found on RBD domain.** The average $\Delta\Delta G_{bind}$ (predicted) is the average of 21 structures.

| **S. No.** | **Mutation** | **RBD-ACE2 interface (Yes/No)** | **Omicron RBD Mutation (Yes/No)** | **Average $\Delta\Delta G_{bind}$ (predicted)** | **Binding (experimental)** | **Expression (experimental)** |
|---|---|---|---|---|---|---|
| 1 | G339D | No | Yes | -0.19 | -0.06 | -0.30 |
| 2 | R346K | No | No | -0.03 | 0.01 | -0.12 |
| 3 | S371L | No | Yes | 0.05 | 0.14 | 0.61 |
| 4 | S373P | No | Yes | 1.04 | 0.08 | 0.22 |
| 5 | S375F | No | Yes | 0.76 | 0.55 | 1.81 |
| 6 | K417N | Yes | Yes | 0.30 | 0.45 | -0.10 |
| 7 | K417T | Yes | No | 0.28 | 0.26 | -0.25 |
| 8 | N439K | No | No | 0.42 | -0.04 | 0.35 |
| 9 | N440K | No | Yes | 0.08 | -0.07 | 0.12 |
| 10 | G446S | Yes | Yes | 1.15 | 0.20 | 0.40 |
| 11 | L452R | No | No | -0.09 | -0.02 | -0.32 |
| 12 | Y453F | Yes | No | 0.35 | -0.25 | 0.08 |
| 13 | S477G | Yes | No | 0.52 | 0.06 | 0.17 |
| 14 | S477N | Yes | Yes | 0.29 | -0.06 | -0.06 |
| 15 | T478K | No | Yes | 0.81 | -0.02 | -0.02 |
| 16 | E484A | Yes | No | -0.01 | 0.07 | 0.23 |
| 17 | E484K | Yes | Yes | 0.09 | -0.06 | 0.10 |
| 18 | E484Q | Yes | No | 0.07 | -0.03 | 0.08 |
| 19 | F490S | Yes | No | 1.09 | 0.00 | 0.10 |
| 20 | Q493R | Yes | Yes | 0.30 | 0.09 | 0.06 |
| 21 | G496S | Yes | Yes | -0.08 | 0.63 | -0.12 |
| 22 | Q498R | Yes | Yes | 0.55 | 0.06 | 0.10 |
| 23 | N501T | Yes | No | 0.15 | -0.10 | 0.25 |
| 24 | N501Y | Yes | Yes | -0.96 | -0.24 | 0.14 |
| 25 | Y505H | Yes | Yes | 1.16 | 0.71 | -0.16 |

**Table S4. Number of RBD mutations with stabilizing, neutral and destabilizing effects.**

| Effect | Number of mutations | Defined range of values |
|---|---|---|
| **$\Delta\Delta G_{bind}$ (predicted)** | | |
| Stabilizing | 325 | < -0.1 |
| Neutral | 389 | -0.1 to 0.1 |
| Destabilizing | 2676 | > 0.1 |
| **Binding (experimental)** | | |
| Stabilizing | 24 | < -0.1 |
| Neutral | 1214 | -0.1 to 0.1 |
| Destabilizing | 2152 | > 0.1 |

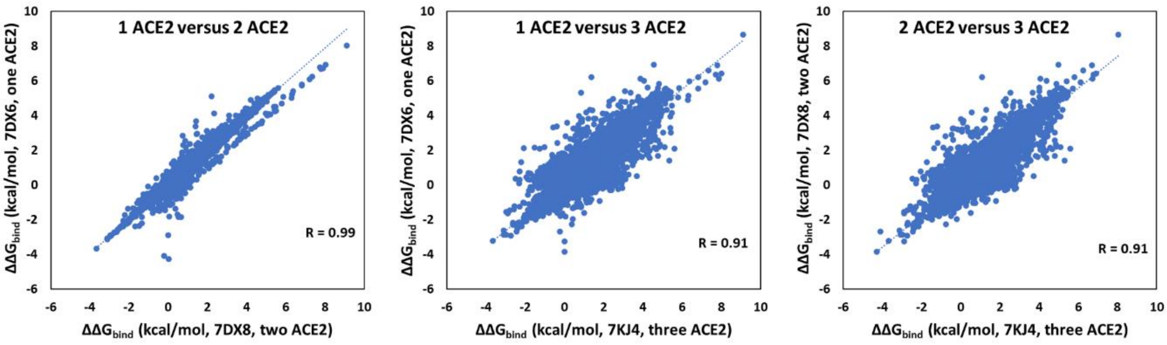

**Figure S1. Comparison of computations using different spike-ACE2 combinations.** Comparison of 1 ACE2 versus 2 ACE2 complexes (7DX6 versus 7DX8), 1 ACE2 versus 3 ACE2 (7DX6 versus 7KJ4), and 2 ACE2 versus 3 ACE2 (7DX8 versus 7KJ4). The PDB structures were selected based on the highest resolutions.

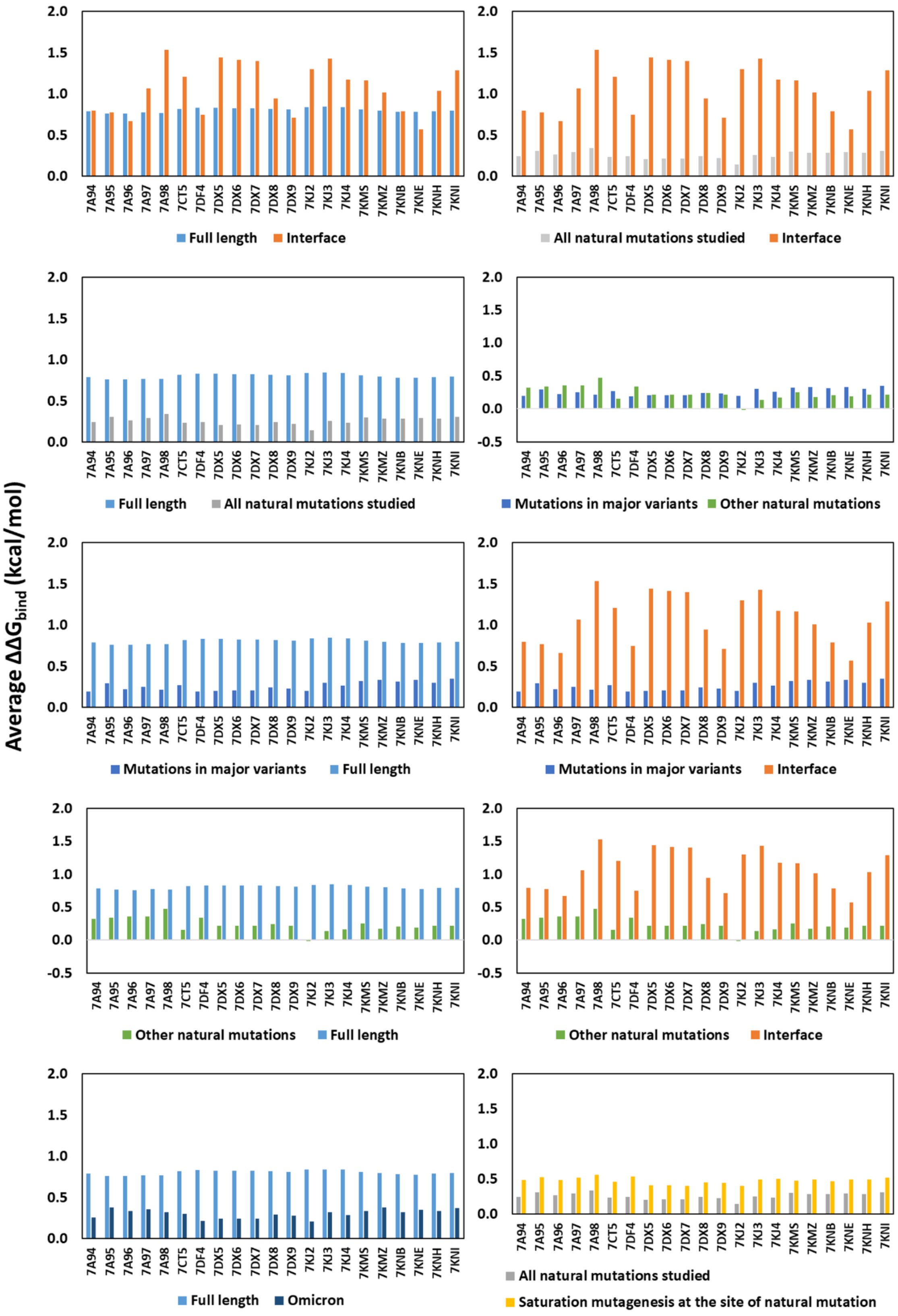

**Figure S2. Comparison of average $\Delta\Delta G_{bind}$ (kcal/mol) for each S-protein-ACE2 complex for different categories of mutations.**

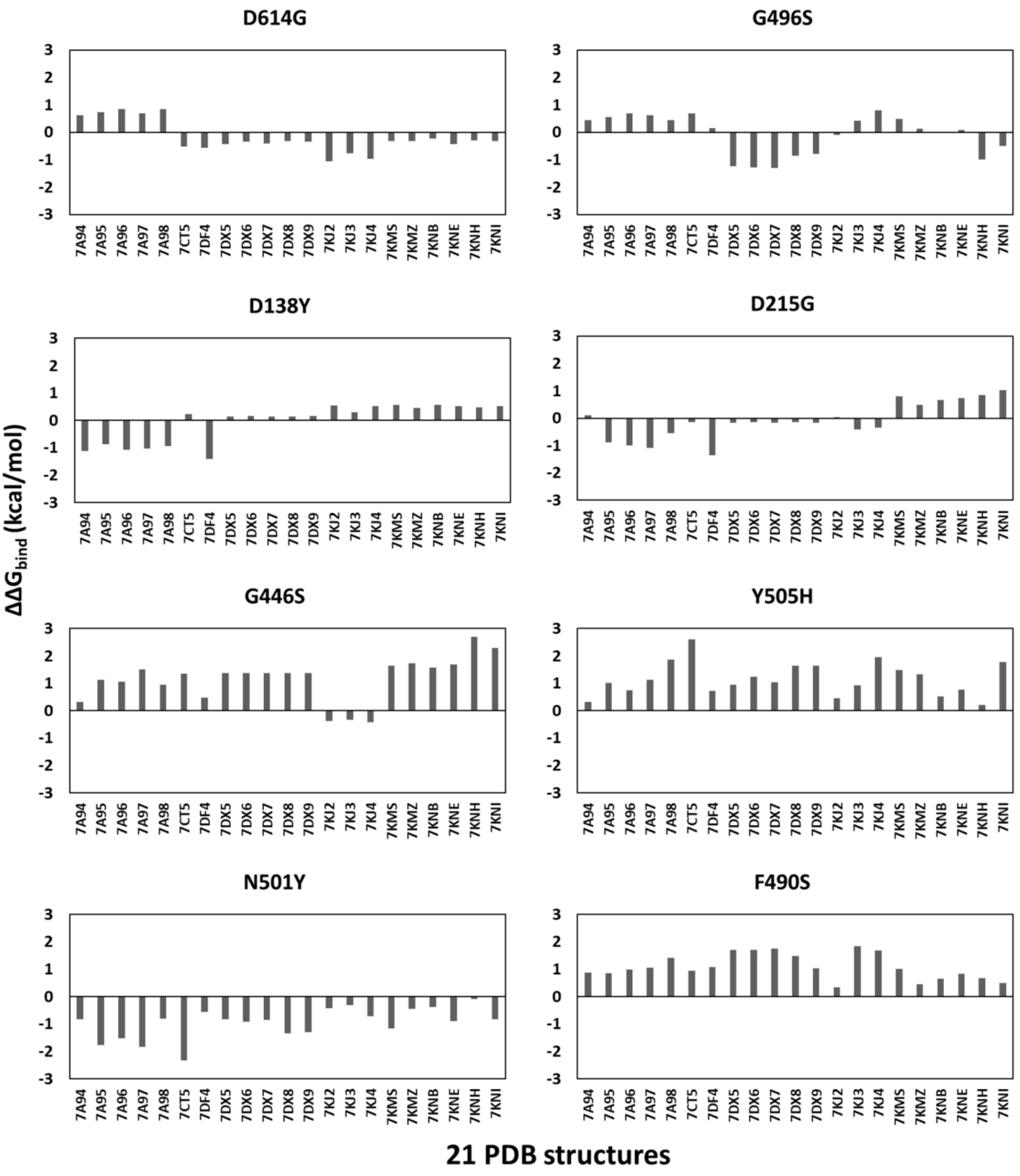

**Figure S3. Variation in the calculated ΔΔGbind (kcal/mol) for several natural mutations calculated using 21 structures showing structural heterogeneity effect on results.**

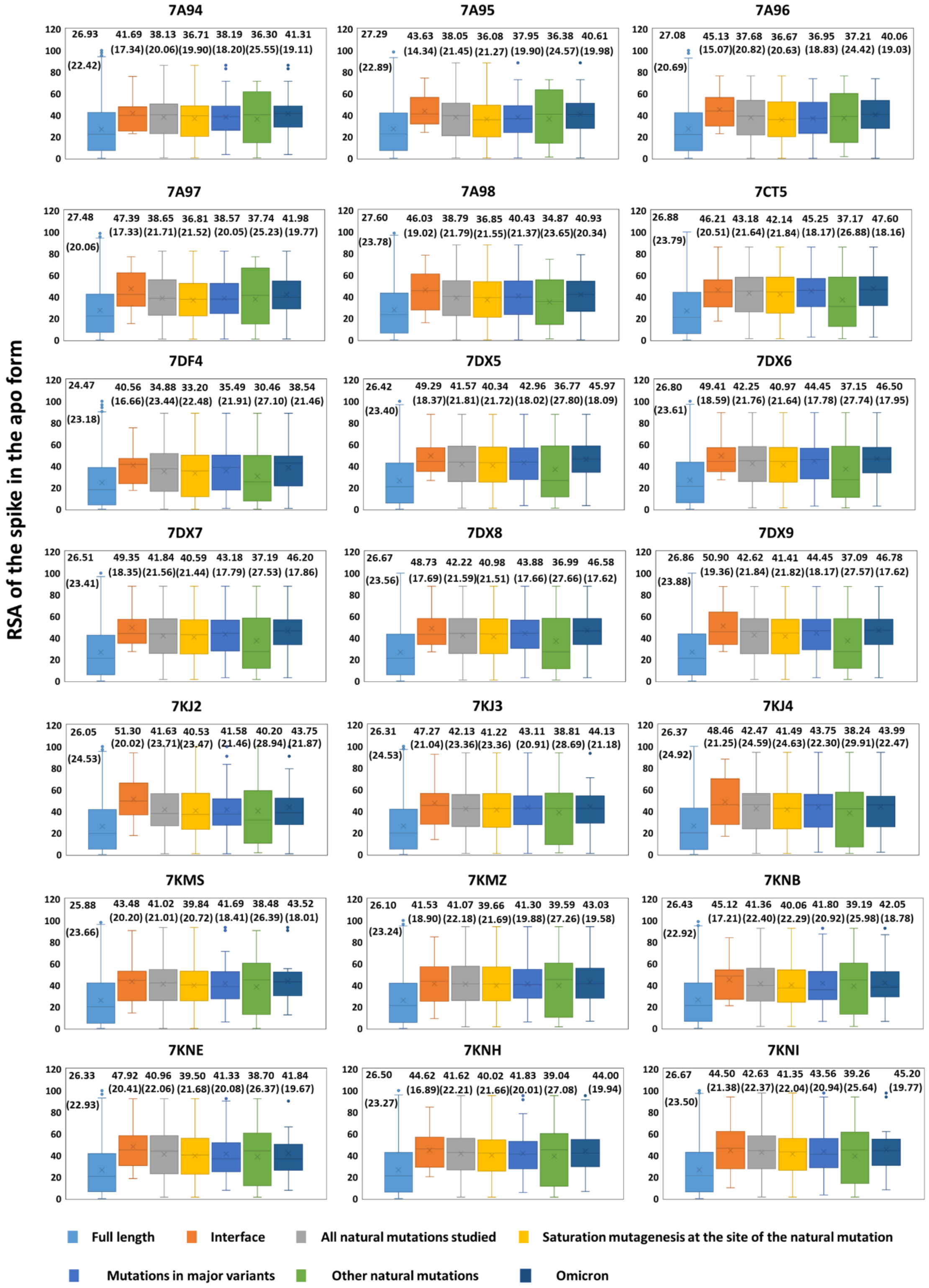

**Figure S4. Residue solvent accessibility for the 21 S-protein structures without ACE2.** The average RSA and standard deviation for each mutation category are indicated in the plots (standard deviation represented inside bracket). The saturation mutagenesis for the site of natural mutations (yellow) is the data averaged over 19 possible mutations for the same site and therefore differ slightly from the dataset of all studied natural mutations (grey).
